# NLRP3 and NLRP1/CARD8 pathways differently contribute to pyroptosis of CD8+ T cells of ART-treated HIV patients

**DOI:** 10.1101/2023.05.15.540712

**Authors:** Mariela EGV Roa, Raylane AG Cambui, Suemy M Yamada, Vinicius CN Leal, Alessandra Pontillo

## Abstract

HIV-infected (HIV) patients exhibit immune dysregulation independently of antiretroviral therapy. The inflammasome, a cytosolic complex responsible for cleavage of the inflammatory cytokines IL -1β and IL -18 and pyroptosis, is highly activated in peripheral blood mononuclear cells of HIV patients, suggesting its involvement in leukocyte dysfunction. While monocytes, B cells, and CD4+ T cells have been studied, little is known about CD8+ T lymphocytes.

Therefore, we proposed to characterize the inflammasome activation in these cells, both the NLRP3 and NLRP1/CARD8 pathways, which are partially described in T cells. CD8+ T lymphocytes from non-HIV healthy donors (HD) and HIV patients were analyzed *ex vivo* and stimulated *in vitro* with known activators of NLRP3 (α-CD3/α-CD28), NLRP1 and CARD8 (DPP9 inhibitor ValboroPro, VbP) to assess inflammasome activation. HIV CD8+ T cells present a constitutively activated caspase-1 which positively correlates with the cell activation state. HIV CD8+ T cells were more activated and more resistant to VbP-induced pyroptosis than HD. On the other way, HIV CD8+ T lymphocytes showed higher pyroptosis in response to α-CD3/α-CD28.

These findings suggest that the NLRP3 pathway is significantly dysregulated in those patients, and TCR stimulation may result in cell loss. At the same time, being HIV CD8+ T cells constitutively activated, other inflammasome pathways, such as NLRP1 or CARD8, present a delayed activation.

## Introduction

Chronic inflammation in patients undergoing antiretroviral therapy (ART) treatment is a consequence of several factors, such as the cytotoxicity of the drugs themselves, viral replication (despite minimal viral reservoirs), damage to the intestinal mucosa (which has been associated with a decreased in the lymphocyte population and regulatory T cells), and, although still debated, increased intestinal permeability, which could lead to microbial translocation from the intestinal lumen to the circulation (Deeks et al., 2015).

The inflammasome, a cytosolic complex responsible for producing IL-1β and IL-18, as well as triggering proinflammatory lytic death known as pyroptosis, appears to be more activated in peripheral blood mononuclear cells (PBMC) of HIV-infected patients, regardless of the use of antiretroviral treatment (Bandera et al., 2018). Furthermore, due to increasing interest in the involvement of the inflammasome in lymphocytes and adaptive immunity, the NLRP3 inflammasome has been described in T lymphocytes, where it induces pyroptosis (Doitsh et al., et al., Monroe et al., et al., 2021) and/or release of IL-1β, at least in CD4+ T cells. In addition, the stimulation of the T cell receptor (TCR) (through CD3 and CD28) induces a not fully characterized NLRP3-dependent caspase-1 activation which involves the production of *r*eactive oxygen species (*ROS*), known up-stream activators of this receptor (Arbore et al., 2016). More recently, it has been demonstrated that chemical inhibition of dipeptidyl peptidase (DPP)-9 by using Val-boroPro (VbP, Talabostat) activates caspase-1-dependent pyroptosis through the engagement of the inflammasome receptors NLRP1/CARD8 in CD4+ T cells (Linder et al., 2021; Johnson et al., 2021).

Since T cells are profoundly disturbed in HIV patients (Fenwick et al., 2019), showing a senescent profile despite ART treatment, and considering that the inflammasome is critical in the development of immunosenescence (Deeks, 2011), it is expected that inflammasome dysregulation contributes to immune dysfunction in HIV patients, in both innate and adaptive immune compartments (Ferrucci and Fabbri, 2018).

Previous studies have demonstrated the crucial roles of inflammasome components in T-cell biology. For example, stimulation of CD3 and CD28 has been shown to induce NLRP3-dependent IL-1β release in CD4+ T cells and to contribute to IFN-γ production and Th1 differentiation (Arbore et al., 2016). In addition, the CARD8 inflammasome has also been identified as a critical regulator of pyroptosis in CD3+ T cells (Linder et al., 2020; Johnson et al., 2020).

In the context of HIV-1 infection, the DNA receptor IFI16 has been found to detect the viral genome and trigger caspase-1-dependent pyroptosis during non-productive HIV infection of CD4+ T lymphocytes (Doitsh et al., 2014; Monroe et al., 2014). More recently, the CARD8 receptor has been identified as the sensor for HIV-1 protease in CD4+ T cells, highlighting the interaction between the inflammasome and HIV-1, not only in innate immune cells but also in lymphocytes (Wang et al., 2021; Clark et al., 2022).

In the past decade, our research group has observed dysregulation of the NLRP3 inflammasome in myeloid cells and B lymphocytes of HIV patients receiving antiretroviral therapy (ART). Interestingly, we found opposite dysfunctions in the innate and adaptive immune compartments. In myeloid cells, the NLRP3 pathway appeared to be unresponsive (Pontillo et al., 2013; Reis et al., 2019; Leal et al., 2020), while in B cells, it was hyper-activated (Leal et al., 2021). Furthermore, recent findings indicate that the inhibition of dipeptidyl peptidase (DPP)-9 can induce pyroptosis in CD4+ T lymphocytes of HIV patients (Lao et al., 2022). This suggests that the NLRP1 and/or CARD8 receptors may also contribute to cell loss and inflammation in chronic HIV infection.

Given the activation/exhaustion of CD8+ T lymphocytes observed in HIV patients (as recently reviewed by Perdomo-Celis et al., 2019) and the possibility of an inflammasome activity in those cells (Arbore et al 2018), we hypothesize that the increased chronic inflammation characteristics of the patients contribute to a dysregulation of the activation of the inflammasome complex also in CD8+ T lymphocytes, as previously observed in other cells.

## Methods

### HIV patients and Healthy donors

All methods were carried out in accordance with the principles of the Declaration of Helsinki and previously approved by the Institute of Biomedical Sciences (ICB) Review Board (CAAE: 52647921.8.0000.5467).

In this study, we recruited ten adult HIV-1–infected (HIV) patients at the “Serviço de Extensão ao Atendimento de Pacientes com HIV/AIDS” of the “Hospital das Clínicas”/Faculty of Medicine, University of São Paulo (HC-FMUSP) (SP, Brazil). All patients are receiving successful combined antiretroviral therapy (ART) for more than 5 years and have had stable CD4+ T cell count (> 500 cells/μL) during the last year (median: 827 cells/μL; I.R.: 735-910 cells/μL) and undetectable viral load (< 50 copies/ml). In addition, patients included in this study do not receive Abacavir (potent activator of NLRP3) (Toksoy et al., 2017), are negative for HBV and HCV, in a generally healthy state (no AIDS, moribund status, active tuberculosis or fungal infections), and without any chronic disease (i.e.: type 2 diabetes, obesity, autoimmunity). Fifteen (n = 15) HIV-negative adult healthy volunteers were included in the study as the control group (healthy donors, HD). Detailed information on these patients are reported in **Supplementary File 1**.

### CD8+ T Cell isolation and treatment

Circulating CD8+ T lymphocytes were isolated from fresh heparin blood by negative selection using magnetic beads (Miltenyi Biotec) from PBMC, previously isolated by Ficoll-Paque gradient (GE Healthcare), and cultured in RPMI-1640 medium (Gibco, Thermofisher Scientific) supplemented with 10% Fetal Bovine Serum (FBS; Gibco, Thermofisher Scientific). Purity of beads-isolated CD8+ T cells was typically > 92% (see **Supplementary File 2** for a representative flow cytometry analysis, FACS). CD8+ T cells were activated in 96-wells culture plates (25 x10^3^ cells/well) coated with anti-CD3 and anti-CD28 (Biolegend) or by the addition of the DPP-8/9 inhibitor Talabostat (Val-boroPro; VbP) (Tocris Biosciences) at the indicated concentrations and times. In some experiments, cells were pretreated with the caspase-1 inhibitor YVAD (Enzo) (1 μM) or with the NLRP3 inhibitor MCC-950 (Invivogen) (10 μM) for 1 hour, detailed dose-response in **Supplementary File 3**. Nigericin (5 μg/mL) was used as a positive control for pyroptosis. Cell viability was monitored by either propidium iodide (ThermoFisher Scientific) or the Zombie Cell Viability Assay (ThermoFisher Scientific) and Annexin-V staining and FACS analysis. Information about antibodies and their concentration are listed in **Supplementary File 4**.

### FLICA analysis of activated caspase-1

Activated caspase-1 was detected by FLICA staining using the FAM-FLICA® Caspase-1 (YVAD) Assay Kit in accordance with the manufacturer’s instructions (Immuno-Chemistry Technologies) with subsequent FACS and Fluorescent microscopy analysis.

### Immunoblotting of GSDMD

CD8+ T cells were plated at a concentration of 2 × 10^6 cells/ml in RPMI containing 10% human serum in 500 μl in a 24-well plate. Cells were washed 1× in PBS and lysed in RIPA lysis buffer containing protease inhibitor (Sigma), centrifugated at 15,000 *g* for 10 min at 4°C, and lysates were transferred to new tubes. Protein concentration was determined by BCA assay (Pierce BCA protein assay kit, Thermo Fisher Scientific) according to the manufacturer’s instructions, and protein amounts were adjusted to 50 ug of protein with 1x Lammli buffer, 95°C for 10 min (Jakobs et al, 2013). Lysate samples were separated by tris-glycine denaturing SDS-PAGE (15% for lysate). Proteins were blotted onto 0.2 mM nitrocellulose membranes (Biorad), blocked in 2.5% BSA (Sigma-Aldrich) in TBS-T, and incubated with the indicated primary (in 2.5% BSA TSBT) and corresponding secondary antibodies (in 2.5% BSA TSBT). Chemiluminescent signals were recorded on ImageQuant LAS 500 (Thermo), and images were analyzed on ImageJ (NIH, [National Institutes of Health](https://www.nih.gov/)).

### Detection of *reactive oxygen species* (*ROS*)

ROS staining was performed by incubating cells with dihydroergotamine 123 (17 mg/ml) diluted in Hank’s balanced salt solution with 10mMHEPES (all from Sigma Aldrich) for 15 min at 37°C. MFI data were measured by fluorimeter (Synergy Biotek Instruments).

### Flow cytometric analysis

Beads-isolated CD8+ T cells were stained extracellularly using primary antibodies for specific detection of surface markers (CD8, CD38, CD107a, TIM3) for 30 minutes at 4°C. For subsequent intracellular staining (IFN-ϒ, caspase-1), cells were permeabilized and fixed using the Permeabilization/Fixation Kit according to the manufacturer’s instructions (eBioscience), and then incubated for 30 minutes at 4°C with primary antibodies (1:50). Samples were acquired by flow cytometry using a BD Canto II flow cytometer (BD Biosciences) and analyzed with FlowJo software (BD Biosciences). At least 10,000 single cells were acquired per sample, with debris and doublets excluded based on their area and aspect ratio.

### Quantification of cytokines

Plasma and culture supernatants were collected and frozen at -80°C until cytokines measurement by commercial ELISA kits (human IL-1ß and IL-18 R&D Systems; human TNF-α and IFN-*γ*, Biolegend).

### Propidium Iodide (PI) real-time incorporation assay

Cells were incubated with Propidium Iodide (PI, Invitrogen, Thermofisher Scientific) (2.5 μg/ml) from 20 to 24 hours post-treatment, and the PI up-take was measured every 5 min on a real-time fluorimeter (Synergy Biotek Instruments) (Case & Roy, 2011; Pierini et al., 2012). The percentage of PI incorporation was calculated based on CD8+ T cells treated with 0.1% Triton X-100 (Sigma Aldrich) for 15 min (= 100 %). As a further correction, the first time point of the kinetics was set to 0.

### ASC “specks” detection and immunofluorescence staining

CD8+ T cells were plated at 0.1 x106 per well of a poly-lysine-treated 96-well chamber slide (Thermo Fisher Scientific), coated with anti-CD3 2.5 μg/mL and anti-CD28 2 μg/mL and activated for 4 hours or stimulated with VbP (5 μM) for 4 hours. Cells were then stained for NLRP3 (Abcam), ASC (Invitrogen), NLRP1 (Abcam), CARD8 (Abcam), or Caspase-1 over-night 4°C and with fluorochrome-conjugated secondary antibody (Thermo Fisher Scientific) for 1 hour. The chamber slide was mounted using the *ProLong Gold Mounting Medium* containing the nuclear dye DAPI (Invitrogen, Thermo Fisher Scientific). Cells were observed using fluorescence microscopy (AxioVert.A1; Zeiss) at 63x magnification with an oil immersion lens. Images were acquired and quantitatively analyzed using ImageJ software (National Institutes of Health, NIH). Five fields for each condition were analyzed for puncta/specks formation according to (Stutz et al. 2013) and for mean fluorescence intensity (MFI). Data were expressed as the number of ASC specks per number of cells per field and for MFI for NLRP1 and CARD8.

### RNA extraction, reverse transcription polymerase chain reaction (RT-PCR), and qPCR

Total RNA was extracted with the *RNAqueous™-Micro Total RNA Isolation Kit* (Ambion, Thermofisher Scientific), and reverse transcription was performed with the Superscript First Strand cDNA Synthesis Kit (Invitrogen, Thermo Fisher Scientific). Relative mRNA expression was examined by qPCR gene-specific Taqman® assays (Applied Biosystems, Thermofisher Scientific) (*NLRP1, NLRP3, CARD8, PYCARD/ASC, CASP1, IL1B, IL18, GSDMD, DPP9)* with a QuantStudio 3.0 real-time PCR equipment (Applied Biosystems, Thermofisher Scientific). Raw expression data (Ct) for each target gene were normalized for the expression of the housekeeping gene *GAPDH* (ΔCt), and the relative expression was shown as 2^-ΔCt (Livak and Schmittgen, 2001).

### Statistical Analysis

Data are presented as mean ± SEM as a pool of at least three experiments, except where it is differently specified. To compare quantitative variables between treated and untreated groups, Student’s paired t-test or one- or two-way analysis of variance (ANOVA) with a Tukey multiple comparison post hoc test or with Sidak multiple-comparisons test, as appropriate. Correlation analysis was performed with Pearson’s correlation test. The threshold for statistical significance was defined at p < 0.05. GraphPad Prism 9 (GraphPad Software) was used for all analyses and graphs realization. Correlation graphs were obtained with the corrplot package and R-project (http://www.R-project.org).

## Results

### CD8+ T cells of ART-treated HIV patients are positive for activated caspase-1 and resistant to TCR-induced pyroptosis

CD8+ T cells isolated from HIV patients present an active exhausted profile compared to healthy donors as assessed by increased expression of surface markers CD38 and TIM-3 (**Figure 1A**) and significantly increased levels of plasma IL-18 and IFN-y when compared to healthy donors (**Fig.1B**). HD and HIV CD8+ T cells did not present a significant difference in IFN-y content or CD107a surface exposure (**Fig.1A**), nor in plasma levels of IL-1ß, which results very low in all samples (**Fig.1B**). These findings confirm that not only CD4+ T lymphocytes but also CD8+ T lymphocytes present signals of exhaustion in ART HIV patients as previously reported (Rallón et al., 2018).

**Fig. 1.**
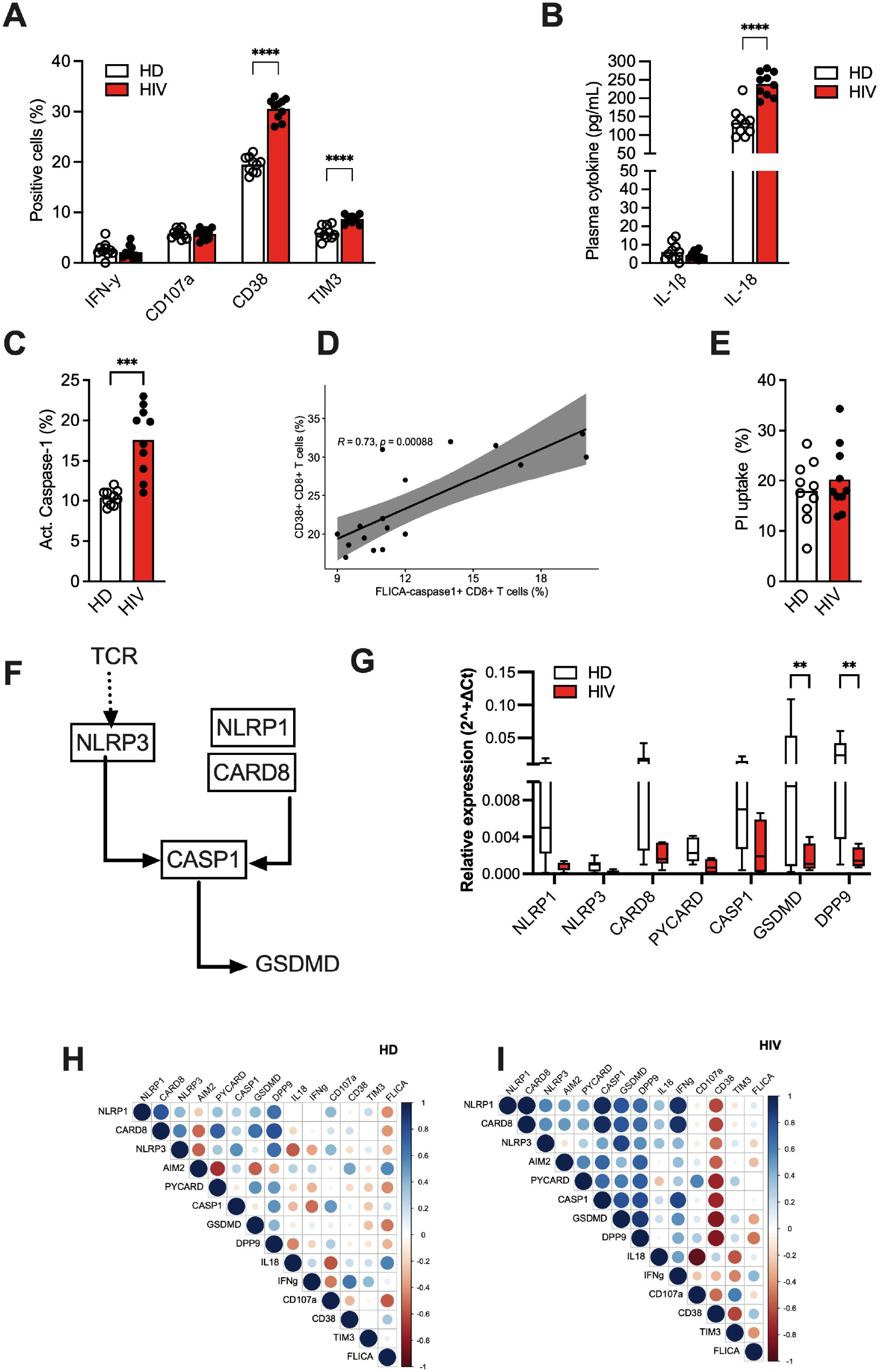
Caspase-1 is activated in CD8+ T cells of HIV patients. Peripheral blood from healthy donors (HD, n = 10) and HIV patients (HIV, n = 10) was used for CD8+ T cells and plasma separation. Flow cytometry analysis of IFN-y and surface markers CD107b, CD38, and TIM-3 was reported as percentage of positive cells (%) for HD and HIV cells **(A)**. Plasma levels of IL-1ß and IL-18 **(B)** were measured in HD and HIV subjects by ELISA. Activated caspase-1 was determined using a FAM-FLICA kit and flow cytometry was reported as a percentage of positive cells (%) (**C)**. Pearson Correlation test was realized for CD38 surface expression and activated caspase-1 (**D)**. The Propidium Iodide (PI) up-take was measured and expressed as the percentage of PI-positive cells related to Triton-treated cells (100%) **(E)**. Relative expression was measured in HD and HIV CD8+ T cells by qPCR and expressed by 2^-ΔCt^ **(G)**. Correlation matrix of inflammasome genes’ expression, plasma IL-18, phenotypic markers, and FLICA– caspase-1 percentage CD8+ T cells in HD **(G)** and HIV patients **(H)**. The size and color density of circles is proportional to the correlation between the 2 variables. The correlation matrix is reordered according to the correlation coefficient using the “hclust” method. A Two-way ANOVA test was applied to compare all the conditions in A, B, and G graphs. The T-test was used in C and E graphs. *: p< 0.05; **: p< 0.01; ***:p< 0.001; ****: p< 0.0001.

PBMC from HIV patients has been shown to present increased caspase-1 activity and ASC-specks formation (Ahmad et al., 2018), suggesting that this activated/exhausted condition may be associated with a dysregulation in the inflammasome complex. In our cohort, a high percentage of HIV CD8+ T cells present caspase-1 activation compared to healthy donors (**Fig.1C**), and a significant positive correlation exists between CD38 surface levels and caspase-1 activation in CD8+ T cells (**Fig.1D**).

We measured either cytokines release or pore-formation by PI uptake assay as possible consequences of inflammasome activation. As expected, no IL-1ß nor IL-18 release was observed in culture supernatants (**data not shown**). Otherwise, no difference in PI uptake was observed (about 20% in HD and HIV samples) (**Fig.1E**).

A reduced expression of inflammasome genes was identified in HIV CD8+ T cells compared to HD (**Fig.1E**), especially *GSDMD* and *DPP9*, reinforcing the hypothesis of a dysregulation of the inflammasome and pyroptosis in HIV lymphocytes.

Finally, while in HD, we cannot observe a significant correlation between CD8+ T cell markers and inflammasome nor with *GSDMD* (**Fig.1F**), in HIV patients (**Fig.1G**), *NLRP1* and *CARD8* expression in CD8+ T cells correlated positively with activated cells (IFN-y+ cells), (p = 0.003, r = 0.923 and p = 0.004, r = 0.916, respectively), as well as with caspase-1 expression (p = 0.001, r = 0.950 and p = 0.011, r = 0.949, respectively), and *GSDMD* (p = 0.038, r = 0.779 and p = 0.042, r = 0.770, respectively). On the other hand, plasma levels of IL-18 negatively correlated with the CD8+ T cell marker CD107a (p = 0.002, r = -0.929). Intriguingly, higher values of correlation were observed between caspase-1, GSDMD, DPP9, and the two receptors NLRP1 and CARD8 just in HIV CD8+ T cells (**Fig.1G**), indicating a basal inflammasome activation and pyroptosis-like signature in HIV CD8+ T cells possibly related to chronic inflammation in those patients, but not in HD (**Fig.1F**).

Kamper’s group has shown that the stimulation of TCR with anti-CD3 and anti-CD28 (α-CD3+α-CD28) induced NLRP3 inflammasome activation and IL-1ß release in CD4+ T cells (Arbore et al., 2016). However, the same condition leads to an unclear response in CD8+ T cells with caspase-1 activation but no cytokine production (Arbore et al., 2018). At that time no data was reported about pyroptosis in those cells, therefore, given the recent findings about pyroptosis in T lymphocytes (Linder et al., 2020: Johnson et al., 2020) and in HIV CD4+ T lymphocytes (Zhang et al., 2021), we evaluated the CD8+ T cells response to the α-CD3+α-CD28 stimulation in term of pyroptosis.

The TCR activation increased the secretion of IFN-*γ* (**Figure 2A**), the surface expression of CD38 and CD107a/LAMP (**Fig.2B-C; Supplementary File 4**) in both HIV and HD CD8+ T cells. On the other hand, the ROS production (**Fig.2D)** and caspase-1 activation (**Fig.2E-F; Supplementary File 5**) were induced only in HIV CD8+ T cells but not in HD.

**Fig. 2.**
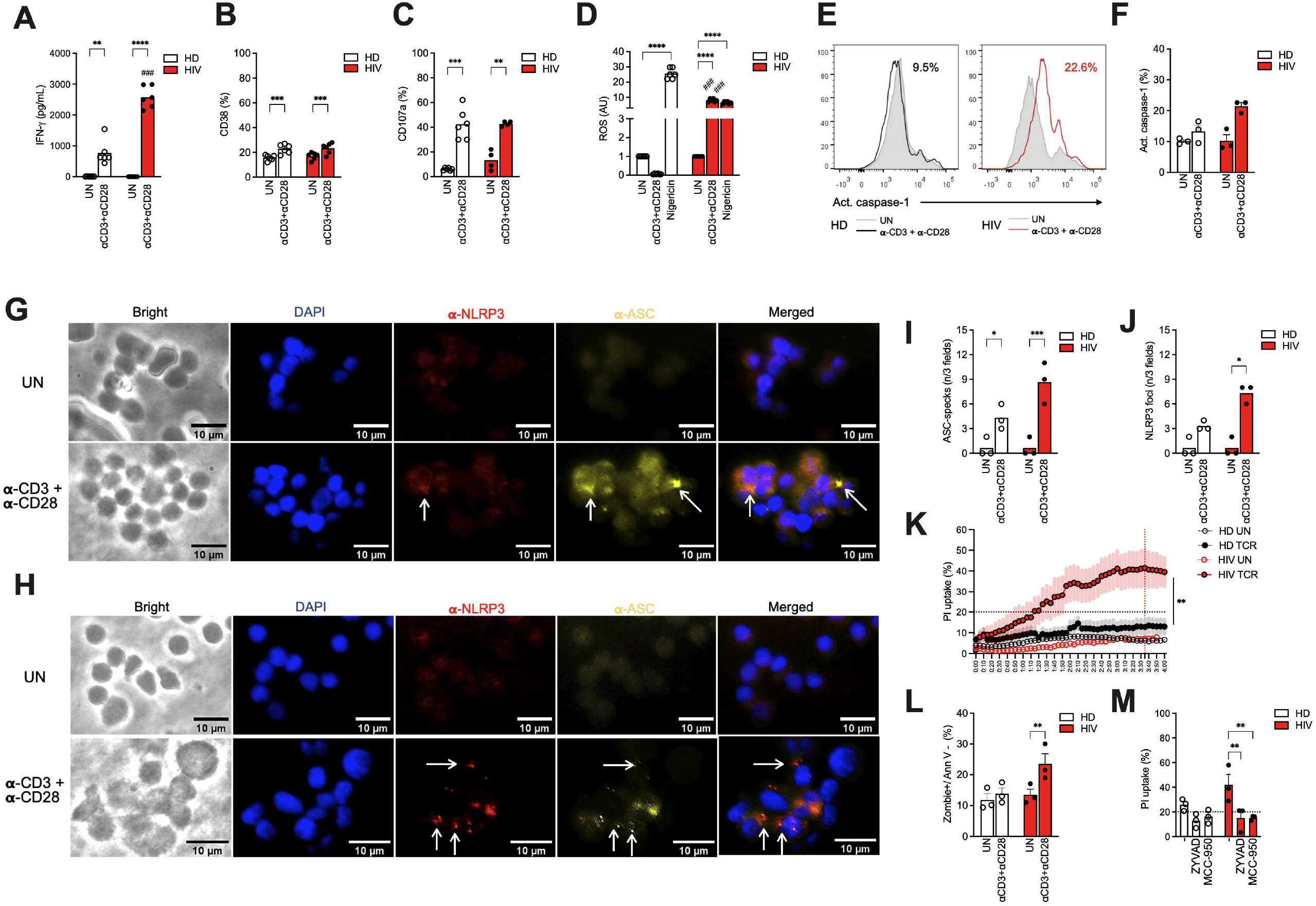
In HIV CD8+ T cells the TCR stimulation triggers NLRP3 inflammasome and pyroptosis. 25.000 CD8+ cells isolated from healthy donors (n = 6) and HIV patients (n = 6) were treated with anti-CD3 (2.5 μg/mL) and anti-CD28 (2 μg/mL) for 48 hours. The release of IFN-ϒ was measured in culture supernatants and expressed as pg/mL **(A)**. The surface expression of CD38 **(B)**, and CD107a/LAMP **(C)**, were analyzed by flow cytometry and expressed as a percentage of positive cells (%). ROS liberation was measured by MitoSOX and measured by Insensitive of fluorescence, and the results were normalized through the basal condition and compared to the other stimulus **(D)**. Activated Caspase-1 was evaluated in CD8+ cells by FAM-FLICA assay and flow cytometry, and expressed as a percentage of positive cells (%). A representative histogram for untreated and treated CD8+ T cells for HD and HIV **(E)** and the average percentages plotted in a bar graph (n = 3) (**F)**. Representative immunofluorescence co-staining for NLRP3 (red) and ASC (yellow) on resting (UN) and αCD3+αCD28 treated CD8+ T cells of HD **(G)** and HIV (**H)** donors. DAPI dye (Blue fluorescence) was used for nuclei coloration. Bar graphs show the average number of ASC specks **(I)** and NLRP3 foci (**J)** in HD and HIV cells (n = 3/group). The Propidium Iodide (PI) uptake was measured and expressed as the percentage of PI-positive cells related to Triton-treated cells (100%) in a real-time curve **(K)**. Cell death (Zombie dye) and Annexin V staining were measured by flow cytometry in resting (UN) and αCD3+αCD28 treated CD8+ T cells of HD and HIV donors (n = 3/group) **(L)**. The effect of caspase-1 inhibitor ZYVAD or NLRP3 inhibitor MCC-950 on PI uptake was measured by real-time assay and the percentage of PI-positive cells are reported HD and HIV cells (n = 3) at 3 hours and 35 minutes of data acquisition **(M)**. Data are reported as average (± standard error). Two Way ANOVA was used to compare the experimental conditions within the two groups.*: p< 0.05; **: p< 0.01; ***: p< 0.001;. ****: p< 0.0001.

Looking at inflammasome mounting by the immunofluorescence staining of ASC-specks and the expected TCR-induced receptor NLRP3 (Arbore et al., 2016), we observed a significant increase of CD8+ T cells positive for ASC-specks in both groups (**Fig.2G-I**), and for NLRP3 oligomerization (*foci)* just in HIV CD8+ T cells (**Fig.2G-H, 2J**). HIV CD8+ T cell morphology seems to be modified by αCD3+αCD28 stimulation (**Fig.2H):** cells are swelled and some bubbles appear, features which may suggest pyroptotic cell death. Therefore, considering that no IL-1ß or IL-18 release was detected in cell supernatants of the two groups (data not shown), we looked forward to the other possible result of inflammasome activation, GSDMD-mediated pores formation.

To evaluate the effect of GSDMD pores formation, we then realized a real-time PI up-take assay to assess the plasma membrane permeability. αCD3+αCD28 stimulation induced a significant increase in PI uptake reaching a maximum after 3 hours 35 minutes of acquisition in HIV CD8+ T cells (41.5 ± 24.6 %), but not in HD ones (12.3 ± 9.1 %) (**Fig.2K**). Of note the majority of TCR-activated HIV CD8+ T cells are doubled-stained for activated caspase-1 and PI (**Supplementary File 6)**, suggesting that the activation of caspase-1 results in pyroptosis. To better characterize the type of cell death, we measured the viability of CD8+ T cells and the expression of apoptosis marker Annexin-V. Flow cytometry data revealed that the αCD3+αCD28 stimulation significantly diminished the viability (increasing Zombie dye incorporation) without increasing Annexin-V expression in HIV cells, but not in HD ones (**Fig.2L, Supplementary File 7**).

These results indicate that HIV CD8+ T cells not only presented increased basal activation of caspase-1 but also that when cells are stimulated, they are more sensitive to NLRP3 inflammasome activation and pyroptosis. To confirm the involvement of caspase-1 and NLRP3 in pyroptosis, we replicated the experiments with chemical inhibitors for caspase-1 (Z-YVAD) or NLRP3 (MCC-950). When CD8+ T cells were pre-treated with the caspase-1 or NLRP3 inhibitor before αCD3+αCD28 stimulation, the PI up-take was dramatically reduced (**Fig.2M**).

Altogether these data show that in HIV patients CD8+ T cells are more sensitive to TCR signaling in terms of inflammasome activation possibly through the NLRP3 receptor, and that this activation leads to caspase-1 activation and pyroptosis.

### Dipeptidyl-peptidase inhibition triggers a lower pyroptosis rate in CD8+ T cells of ART-treated HIV patients compared to healthy donors

Recently the DPP-inhibitor Val-boroPro (VbP) has been described to induce pyroptosis in human T cells (Linder et al., 2020: Johnson et al., 2020), even if the effect on inflammasome activation specifically in CD8+ T cells have not been completely elucidated.

VbP (5 μM) (see **Supplementary File 8** for time and dose-response tests) significantly increased the surface expression of CD38 (**Figure 3A**) and CD107a/LAMP (**Fig.3B**) in HIV CD8+ T lymphocytes, but it did not induce significant secretion of IFN-γ, IL-1ß or IL-18 (above detection limits; data not shown). Caspase-1 activation was observed in both HD and HIV VbP-treated CD8+ T cells, even if the effect was significantly lower in HIV than in HD cells (**Fig.3C-D**). Accordingly, VbP induced a significant increase in membrane permeability in CD8+ T cells in both groups. However, the PI up-take was lower in HIV than in HD (**Fig.3E-F; Supplementary File 9**). In addition, VbP induced Annexin V-negative cell death in HD and HIV lymphocytes, however, just in HD cells the increment was statistically significant (**Fig.3G)**. The VbP-induced PI up-take is dependent on caspase-1 activity, and independent from NLRP3, as demonstrated by the pre-treatment of CD8+ T cells with the two inhibitors ZYVAD and MCC-950 (**Fig.3H)**.

**Fig. 3.**
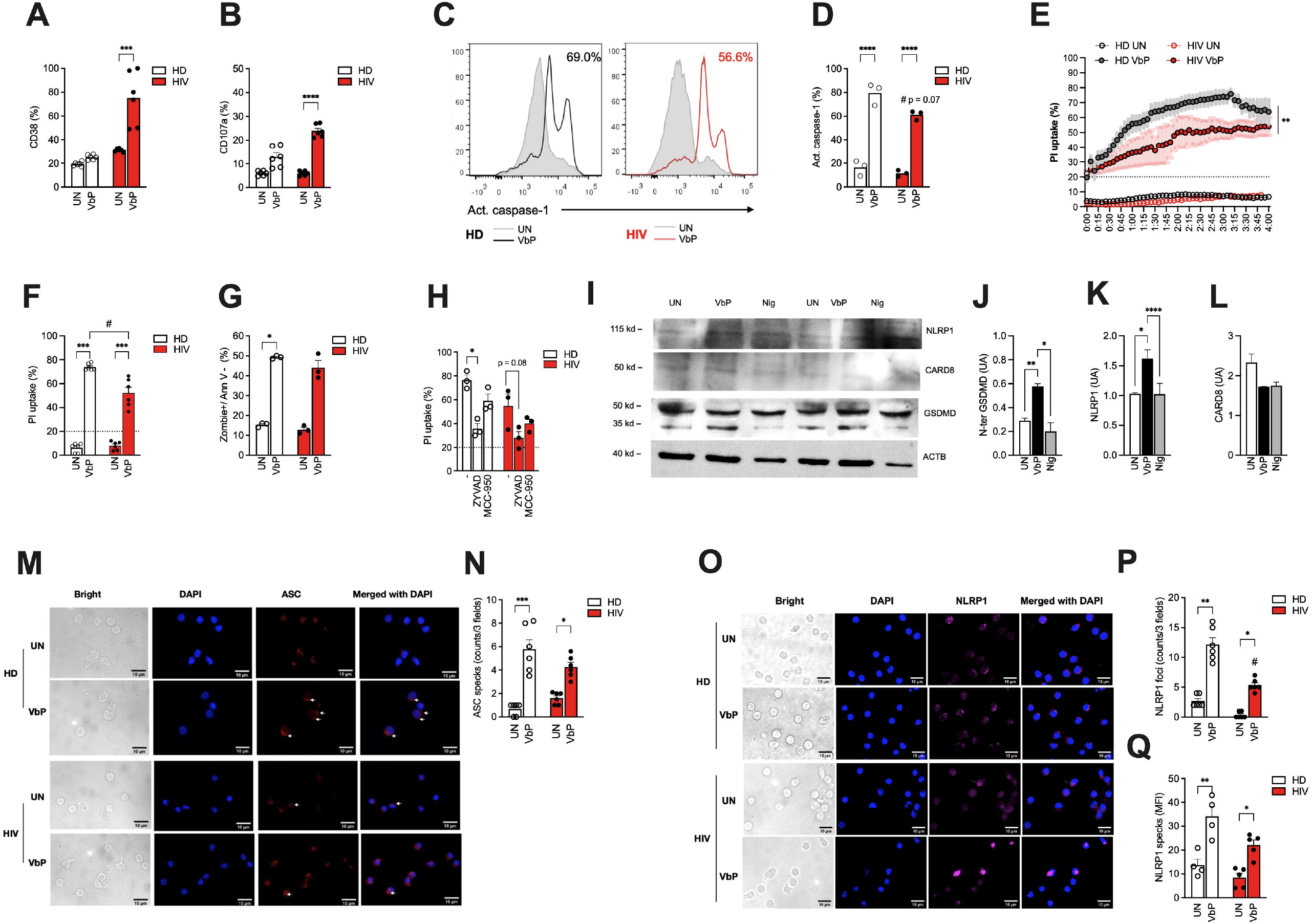
The DPP-9 inhibition activates caspase-1 and pyroptosis in human CD8+ T cells. 25.000 CD8+ cells isolated from HIV patients (HIV; n = 6) and healthy donors (HD; n = 6) and treated with Valboro-Pro (VbP; 5 μM) for 24 hours. The surface expression of CD38 **(A)** CD107a/LAMP **(B)** and caspase-1 cleavage/activation **(C-D)** were analyzed by flow cytometry and expressed as a percentage of positive cells (%). The Propidium Iodide (PI) uptake was measured and expressed as the percentage of PI-positive cells related to Triton-treated cells (100%) in a real-time curve **(E)** and as an end-point bar graph at 3 hours and 35 minutes of incorporation **(F)**. The Annexin V-negative and Zombie-positive cell death was measured by flow cytometry and expressed as a percentage of positive cells (%) (n = 3) **(G)**. The effect of caspase-1 inhibitor ZYVAD or NLRP3 inhibitor MCC-950 on PI uptake was measured by real-time assay and the percentage of PI-positive cells are reported at 3 hours and 35 minutes of incorporation (n = 3) at 3 hours and 35 minutes of data acquisition **(H)**. Representative immunoblot for GSDMD, NLRP1, CARD8, and endogenous GAPDH in untreated and VbP-treated CD8+ T cells (lysed cells) of HD and HIV subjects (n = 2). Treatment with Nigericin 5 μM was used as a positive control for GSDMD pores formation **(I)**. Mean luminescence intensity for GSDMD (J), NLRP1 (K), and CARD8 (L) proteins relative to the endogenous GAPDH. Representative immunofluorescence staining of ASC (M) and NLRP1 (O) in untreated and VbP-treated CD8+ T cells of HD and HIV subjects. The number of ASC specks **(N)** or NLRP1 foci/cell **(P)** and mean fluorescence intensity (MFI) for NLRP1 **(Q)** are reported for untreated and VbP-treated CD8+ T cells of HD and HIV subjects (n = 3). Data are reported as average (± standard error). One or Two Way ANOVA was used to compare the experimental conditions within the two groups (*) and between the groups (#).*: p< 0.05; **: p< 0.01; ***: p< 0.001;. ****: p< 0.0001.

These findings point to a VbP-induced pyroptosis, which was confirmed by the increased cleavage of GSDMD (**Fig. 3I-J)**.

Given that the inflammasome receptors NLRP1 and CARD8 are known targets of VbP (Zhong et al., 2018), we examined these two receptors in CD8+ T cells. While both receptors are present in CD8+ T cells, only NLRP1 appears to show a slight increase in VbP-treated cells **(Fig.3I, Fig.3K)**, forming *foci* indicative of complex oligomerization, which is lower in HIV cells compared to HD cells **(Fig.3O-Q)**. This is not the case for CARD8, as its fluorescence appears to increase after VbP treatment **(Fig.3I, Fig.3L)**; however, no foci or oligomerization were observed **(Supplementary File 10)**. Of note, ASC specks are present in VbP-treated CD8+ T cells **(Fig. 3M-N)**, with a significantly greater number observed in healthy donors compared to HIV cells.

These findings showed that the DPP8/9 inhibition induced caspase-1-dependent pyroptosis in CD8+ T cells and that HIV CD8+ T cells were more resistant to VbP-induced pyroptosis than non-HIV healthy donors. According to our findings, NLRP1 may be the receptor responsible for the activation of the inflammasome by VbP in CD8+ T cells.

Given that lymphocytes from HIV individuals are known to be constitutively activated (**Fig. 1**) (Ahamad et al., 2002), and that TCR-activated CD3 lymphocytes have been shown to exhibit less pyroptosis compared to resting cells (Linder et al., 2020), we can therefore attribute the observed resistance to VbP-induced pyroptosis in HIV CD8+ T cells to their activated state. Furthermore, as a proof of concept, when we started CD8+ T cells from healthy donors with αCD3+αCD28 or PMA before VbP treatment, we observed a reduction in VbP-induced pyroptosis and NLRP1 oligomerization (but not CARD8) (**Supplementary File 11**).

## Discussion

Previous research has shown increased expression of activation/exhaustion markers and caspase-1 activation suggestive of constitutive inflammasome assembling in circulating leukocytes from HIV patients (Bandeira et al., 2018; Ahmad et al., 2018). Furthermore, the dysregulation of the inflammasome, and specifically the NLRP3 inflammasome, has been commonly observed in monocytes and monocytes-derived cells in HIV patients as a result of the proinflammatory *milieu* of those patients (Reis et al., 2019). Since the assembly of the NLRP3 inflammasome has been reported in lymphocytes as well (Arbore et al., 2016 and 2018; Linder et al., 2020; Johnson et al., 2020), it has been postulated that similar dysregulation of the inflammasome may impact the threshold or intensity of inflammasome activation in HIV lymphocytes, thereby influencing their biology. Accordingly, CD4+ T lymphocytes in HIV patients exhibit high NLRP3-dependent caspase-1 activation and pyroptosis, which contribute to the depletion of CD4+ T cells during chronic infection, regardless of the use of ART (Zhang et al., 2021). Additionally, the expression of pyroptosis-related genes is elevated in PBMC of ART-treated HIV patients, especially in those categorized as immunological non-responders (Lao et al., 2022). These findings provide clear evidence that the inflammasome and caspase-1-induced pyroptosis play significant roles in chronic inflammation within the lymphoid compartment of HIV patients. However, there is still limited knowledge regarding CD8+ T cells in HIV patients, including their exhausted functionality and their specific contribution to chronic inflammation in HIV infection.

In this study, we conducted experiments to investigate the response of HIV CD8+ T cells to T-cell receptor (TCR) stimulation and DPP-8/9 inhibition. Our findings revealed that the TCR stimulation of HIV CD8+ T cells led to a general activation of the lymphocytes, including increased secretion of IFN-ϒ, and to the activation of the NLRP3 inflammasome. Interestingly, we observed that this activation did not result in the release of the cytokines IL-1β or IL-18, but in an increased pyroptosis. These results suggest that the NLRP3 inflammasome pathway in HIV CD8+ T cells may regulate cell death processes rather than cytokine release. Arbore et al. (2018) recently reported the activation of the NLRP3 inflammasome in CD8+ T cells. However, they did not observe the release of cytokines or the induction of pyroptosis following NLRP3 activation in these cells. Our results showed increased pyroptosis in response to TCR stimulation and elucidated the outcome (or one of the possible outcomes) of NLRP3 inflammasome activation in CD8+ T cells in chronic HIV infection. Indeed, reactive oxygen species have been implicated as upstream factors that can induce NLRP3 inflammasome activation and previous studies (Arbore et al., in 2016 and 2018) have suggested that TCR stimulation can lead to the production of ROS, which in turn may contribute to the activation of the NLRP3 inflammasome in T cells. The release of ROS in HIV lymphocytes is notoriously augmented (Perrin et al., 2012; Zhang et al., 2021; Deguit et al., 2019), and as we demonstrated here, HIV CD8+ T cells produce more ROS than healthy donors, which can result in the up-regulation of NLRP3 inflammasome activation and consequent increment of pyroptosis. Another possible explanation is the increased degranulation in HIV CD8+ T cells, as measured by CD107a/LAMP1. The subsequent release of cytotoxic proteins and the formation of cell membrane pores can lead to an ionic imbalance and potassium efflux, which may activate the NLRP3 inflammasome. However, further research is needed to fully understand the specific mechanisms and interactions involved in NLRP3 activation in HIV CD8+ T cells.

When we challenged CD8+ T cells with the chemical activator of NLRP1/CARD8 inflammasome, Valboro-Pro (Talabostat), we observed the increased activation of caspase-1 followed by pyroptosis, but not IL-1ß/IL-18 release, in healthy donors and HIV patients. Therefore suggesting that this pathway for inflammasome activation is present in CD8+ T cells as it is in CD4+ T cells (Linder et al., 2021; Johnson et al., 2021). Unexpectantly VbP induced the increased expression of activation markers CD38 and CD107a just in HIV cells. However, since no IFN-ϒ release was noticed, this effect remains not fully elucidated, even if some works have suggested that VbP increases the immune response in the tumor environment (Henderson et al., 2021; Yang et al. 2023).

As expected, VbP-induced pyroptosis is caspase-1 dependent and NLRP3-independent in HD and HIV CD8+ T cells, however at the moment we cannot be able to distinguish between NLRP1 and CARD8, as possible targets of VbP. Even though Linder et al. (2021) have previously demonstrated that CARD8 is responsible for VbP-induced pyroptosis in T cells, we are inclined to say that, at least in CD8+ T cells, NLRP1 seems to be more critical than CARD8 in responding to VbP. The immunofluorescence assays clearly showed the visible oligomerization of NLRP1, but not of CARD8, in response to VbP treatment. This oligomerization is compatible with the ASC-specks formation, and up-to-now we know that NLRP1 but not CARD8 recruits the caspase-1 through the adaptor molecule ASC (Ball et al., 2019).

A tricky question is what could be the natural activator of NLRP1 (or CARD8) in CD8+ T cells. To our knowledge both receptors are demonstrated to be targets of viral proteases (Tsu et al., 2021; Nadkarni et al., 2021) inclusive of HIV-1 (Wang et al., 2021); however, CD8+ T cells are not known to be significantly infected by specific viruses. On the other hand, it is still discussed whether cell stress, namely ER stress (D’Osualdo et al 2015; Cao et al., 2019), reductive stress (Wang et al., 2023), or protein folding stress (Orth et al., 2023), or even senescence it-self (Muela-Zarzuela et al., 2023) can activate NLRP1 and/or CARD8 inflammasome.

Finally, we observed a lower rate of pyroptosis in HIV CD8+ T lymphocytes compared to HD. Since Linder et al (2021) have demonstrated that TCR-activated T cells are resistant to VbP-induced pyroptosis, this finding may be a consequence of the constitutive basal activation of CD8+ T cells in HIV patients (Bandeira et al., 2018; Ahmad et al., 2018). Indeed, when we treated HD CD8+ T cells with αCD3+αCD28 and then with VbP, we confirmed the resistance to VbP-induced pyroptosis in these cells, as it was previously demonstrated for whole CD3+ T cells (Linder et al., 2021), and supporting the idea that the resistance of HIV CD8+ T cells to VbP may be the consequence of their activation state.

In conclusion, this study reveals that TCR-induced NLRP3/caspase-1 activation in CD8+ T cells leads to pyroptosis in individuals with HIV, but not in healthy donors. Additionally, we have demonstrated, for the first time to our knowledge, that the NLRP1/CARD8 pathway is present in CD8+ T cells and leads to pyroptosis. Due to the constitutive activation of HIV lymphocytes, HIV CD8+ T cells show resistance to VbP-induced pyroptosis. Finally, our data suggest that NLRP1 is the primary target of VbP, although further investigations are needed to thoroughly explore this pathway in CD8+ T cells.

## Supporting information

https://drive.google.com/drive/folders/1bJB7Ww7-LIlrADNnCu_0ycJ9fg1FEaC9?usp=share_link

