## Supplementary material for "NLRP3 and NLRP1/CARD8 pathways differently contribute to pyroptosis of CD8+ T cells of ART-treated HIV patients": https://drive.google.com/drive/folders/1bJB7Ww7-LIlrADNnCu_0ycJ9fg1FEaC9?usp=share_link

Supplementary File 1 Characteristics of the volunteers recruited for the Study

| **ID** | **Sex** | **Age, year** | **Disease time, years** | **CD4+ T (cells/µL)** | **CD8+ T (cells/µL)** | **CD4/CD8 ratio** | **ART time, years** |
| --- | --- | --- | --- | --- | --- | --- | --- |
| HIV01 | F | 25 | 16 | 1765 | 1548 | 1.14 | 16 |
| HIV02 | M | 71 | 15 | 823 | 701 | 1.17 | 14 |
| HIV03 | F | 64 | 9 | 621 | 523 | 1.19 | 8 |
| HIV04 | M | 62 | 26 | 978 | 613 | 1.60 | 25 |
| HIV05 | M | 64 | 27 | 956 | 650 | 1.47 | 26 |
| HIV06 | M | 60 | 25 | 1237 | 808 | 1.53 | 25 |
| HIV07 | F | 62 | 23 | 749 | 595 | 1.26 | 23 |
| HIV08 | F | 62 | 25 | 789 | 543 | 1.45 | 24 |
| HIV09 | M | 65 | 22 | 937 | 725 | 1.29 | 21 |
| HIV10 | M | 57 | 21 | 740 | 545 | 1.36 | 20 |
| HD01 | F | 49 |  |  |  |  |  |
| HD02 | M | 29 |  |  |  |  |  |
| HD03 | F | 30 |  |  |  |  |  |
| HD04 | F | 32 |  |  |  |  |  |
| HD05 | F | 27 |  |  |  |  |  |
| HD06 | F | 28 |  |  |  |  |  |
| HD07 | M | 48 |  |  |  |  |  |
| HD08 | M | 28 |  |  |  |  |  |
| HD09 | F | 29 |  |  |  |  |  |
| HD10 | M | 24 |  |  |  |  |  |
| HD11 | M | 28 |  |  |  |  |  |
| HD12 | F | 48 |  |  |  |  |  |
| HD13 | F | 20 |  |  |  |  |  |
| HD14 | F | 50 |  |  |  |  |  |
| HD15 | M | 29 |  |  |  |  |  |

F, Female. M, Male. SD..

Supplementary File 2. Purity and viability of beads-isolated CD8+ T cells

The strategy of gates and dot-blots was shown for a representative experiment.

Briefly, 0.5 x10^6^ CD8+ T cells isolated from a healthy donor were stained for CD8 surface marker and analyzed by flow cytometry.

### Supplementary File 3. Inhibitors dose-response

Dose-response of the NLRP3 and Caspase-1-4 inhibitors was performed to understand the best work concentration, adapted for our number and concentration of cells. 25.000 CD8+ cells isolated from healthy donors (n = 3) and were treated with several concentrations of MCC950 (**A**) and ZYVAD (**B**) previously 30 minutes before the 5 µM VbP for 24 hours, and 2.5 µg/mL of anti-CD3 and 2 µg/mL of anti-CD28 for 48 hours.

Supplementary File 4.  Antibodies e reagents

| **ANTIBODY** | **SOURCE** | **IDENTIFIER Catalog (C#) or Reference (R#)** | **CLONE** | **DILUTION % (v/v)** |
| --- | --- | --- | --- | --- |
| Anerxin V - FITC | BD | C#. 556420 | DX2 | 1 to 50 |
| anti-CD28 | Biolegend | C#. 302904 | CD28.2 | 2 µg/mL |
| anti-CD3 | Biolegend | C#. 300438 | UCHT1 | 2.5 µg/mL |
| ASC - Mouse | Merck | C#. 04147 | 2EI7 | 1 to 200 |
| CARD8 - rabbit | Abcam | ab24186 | Polyclonal | 1 to 200 |
| Caspase-1 - rabbit | Abcam | ab626988 | Polyclonal | 1 to 200 |
| CD107a - Pacific Blue | Biolegend | C#. 328620 | H4A3 | 1 to 60 |
| CD38 - APC | Beckman Coulter | R#. IM1832U | T16 | 1 to 40 |
| CD69 - APC-Cy7 | BD | C#. 557756 | FN50 | 1 to 40 |
| CD8 - PE | Biolegend | C#. 344706 | SK1 | 1 to 50 |
| Gasdermin D - Rabbit | Cell signaling | C#. 39754 | E9S1X | 1 to 1000 |
| Goat- anti- Rabbit - HRP | Abcam | ab97080 | Polyclonal | 1 to 5000 |
| Goat-anti-Mouse - AF488 | Abcam | Ab181347 | Polyclonal | 1 to 600 |
| Goat-anti-Rabbit - AF647 | Abcam | ab17547 | Polyclonal | 1 to 1000 |
| NLRP1 - rabbit | Abcam | ab3683 | Polyclonal | 1 to 200 |
| NLRP3 - rabbit | Abcam | ab214185 | Polyclonal | 1 to 200 |
| TIM3- BB700 | Fisher Scientific | C#. BDB746178 | 7D3 | 1 to 50 |

Supplementary File 5 - FACS analysis of TCR-activated CD8+ T cells

Dot-blots were shown for a representative experiment.

Briefly, 0.5 x106 CD8+ T cells isolated from one healthy donor and one HIV patient were stained for CD8 together with CD38 (**A**), CD107a (**B**), and activated caspase 1 (**C**) and analyzed by flow cytometry.

Supplementary File 6 - FAM-FLICA and PI staining in TCR-activated CD8+ T cells

Images were shown for a representative experiment.

Briefly, 0.1 x10^6^ CD8+ T cells isolated from one healthy donor and one HIV patient were treated with a-CD3 and a-CD28 for 48 hours and stained for activated caspase-1 (FAM-FLICA), PI (2.5 µg/mL) and Hoechst dye and observed in fluorescent microscopy.

Supplementary File 7 - FAM-FLICA and PI staining in TCR-activated CD8+ T cells

Images were shown for a representative experiment.

Briefly, 0.1 x10^6^ CD8+ T cells isolated from one healthy donor and one HIV patient were treated with anti-CD3 (2,5µ/mL) and anti-CD28 (2 µg/mL) 24 hours and stained for activated caspase-1 (FAM-FLICA), PI (2.5 µg/mL) and Hoechst dye and observed in fluorescent microscopy.

Supplementary File 8 - Dose Response of VbP-induced Pyroptosis

25.000 CD8+ cells isolated from healthy donors (n = 3) and HIV patients (n = 3) were treated with f 2.5, 5, and 10 µM VbP for 24 hours. The Propidium Iodide (PI) up-take was measured and expressed as the percentage of PI-positive cells related to Triton-treated cells (100%) in a real-time curve (A).

Two-way ANOVA test was applied to compare all the conditions in the below graphs. Associations were evaluated using Spearman’s correlation test *: p< 0.05; **: p< 0.01; ***:p< 0.001; ****: p< 0.0001.

Supplementary File 9 - FAM-FLICA and PI staining in VbP-activated CD8+ T cells

Images were shown for a representative experiment.

Briefly, 0.1 x10^6^ CD8+ T cells isolated from one healthy donor and one HIV patient were treated with VbP 5 µM for 24 hours and stained for activated caspase-1 (FAM-FLICA), PI (2.5 µg/mL) and Hoechst dye and observed in fluorescent microscopy.

Suplementary File 10. The activation of CD8+ T lymphocyte prevent VbP-induced pyroptosis

25.000 CD8+ cells isolated from healthy donors (n = 3) were treated with anti-CD3 (2.5 g/mL) and anti-CD28 (2 µg/mL) for 24 hours and plus 5 uM VbP for 24 hours.  The Propidium Iodide (PI) up-take was measured and expressed as the percentage of PI-positive cells related to Triton-treated cells (100%) in a real-time curve **(A).** NLRP1 agglomeration is stained with red fluorescence (Goat anti-human NLRP3). DAPI dye (Blue fluorescence) was used for nuclei coloration. Data were reported as microscopy photographs (10 µm) for one representative experiment **(D)** and bar graphs showing the MFI and ‘speaks’ or protein agglomerate formation in CD8+ T cells as a mean of the three experiments ± standard error **(B-C).**

### Supplementary File 11. VbP did not induce CARD8 oligomerization in CD8+ T lymphocytes

0.1 x10^6 CD8+ T cells were treated with VbP 5 µM for 24 hours, stained for CARD8 (red fluorescence), and observed in fluorescent microscopy. DAPI dye (Blue fluorescence) was used for nuclei coloration. Data were reported as microscopy photographs (10 µm) for one representative experiment (A), and as bar graphs as CARD8 foci/cell **(B)** and mean fluorescence intensity (MFI) **(C)** are reported for untreated and VbP-treated CD8+ T cells of HD and HIV subjects (n = 3).
